# Molecular pathway of influenza pan-neuraminidase inhibitors resistance in an immunocompromised patient

**DOI:** 10.1101/768614

**Authors:** Yacine Abed, Manuel Schibler, Liva Checkmahomed, Julie Carbonneau, Marie-Christine Venable, Federica Giannotti, Ana Rita Goncalves, Laurent Kaiser, Guy Boivin

## Abstract

Neuraminidase (NA) inhibitors (NAIs), including oseltamivir and zanamivir, play an important therapeutic role against influenza infections in immunocompromised patients. In such settings, however, NAI therapy may lead to the emergence of resistance involving mutations within the influenza surface genes. The aim of this study was to investigate the evolution of hemagglutinin (HA) and NA genes of influenza A(H1N1)pdm09 virus in an immunocompromised patient receiving oseltamivir then zanamivir therapies. Nasopharyngeal swabs (NPS) samples were collected between 01-27-2018 and 04-20-2018 from a hematopoietic stem cell transplant recipient. These included 11 samples collected either pre-therapy, during oseltamivir and zanamivir as well as after therapy. The A(H1N1)pdm09 HA/NA genes were sequenced. The H275Y NA substitution was quantified by droplet digital RT-PCR assay. A(H1N1)pdm09 recombinant viruses containing HA mutations were tested by HA elution experiments to investigate *in vitro* binding properties. Oseltamivir rapidly induced the H275Y NA mutation which constituted 98.33% of the viral population after 15 days of oseltamivir treatment. The related HA gene contained S135A and P183S substitutions within the receptor-binding site. After a switch to zanamivir, 275H/Y and 119E/G/D mixed populations were detected. In the last samples, the double H275Y-E119G NA variant dominated with S135A and P183S HA substitutions. This report confirms that oseltamivir can rapidly induce the emergence of the H275Y substitution in A(H1N1)pdm09 viruses and subsequent switch to zanamivir can lead to additional substitutions at codon E119 resulting in multi-drug resistance. Such data highlight the need for novel antiviral agents.

## 1. Introduction

Influenza viruses are responsible for acute respiratory diseases that constitute a significant public health priority worldwide. During typical seasonal epidemics, influenza viruses affect 10% to 20% of the human population [1]. In most people, seasonal influenza strains replicate in the upper respiratory tract causing a self-limited disease. However, in high-risk individuals, such as immunocompromised patients, viral replication may progress towards the lower respiratory tract, potentially triggering secondary bacterial infections and pneumonia and leading to more severe infections and sometimes death [2]. Hematopoetic stem cell transplant (HSCT) recipients are among populations at higher risk of developing seasonal influenza infections with fatal outcomes [3]. Since their appearance in 2009, influenza A(H1N1)pdm09 viruses continue to circulate worldwide and have become important seasonal respiratory pathogens. A recent report showed that viral shedding in HSCT patients with A(H1N1)pdm09 infections was more prolonged compared to infections with other seasonal influenza A subtypes and B strains [4]. The use of antivirals in this situation has the potential to increase the risk of developing drug resistance.

Anti-influenza agents are expected to play a major role against severe influenza infections. Before the recent approval of baloxavir marboxil in 2018 [5], neuraminidase inhibitors (NAIs) constituted the only class of antivirals recommended for the control of influenza infections. Since their development in late 1990s, oseltamivir and, to a lesser extent, zanamivir have been the most frequently used NAIs worldwide [6]. With the advent of the 2009 influenza A(H1N1) pandemic, the use of oseltamivir has grown globally for both prophylactic and therapeutic purposes increasing the potential for the emergence of NAI resistance [7, 8]. In immunocompromised patients, A(H1N1)pdm09 variants harboring the H275Y NA substitution conferring oseltamivir resistance emerged at rates that could exceed 50% [9, 10]. As A(H1N1)pdm09-H275Y variants retain susceptibility to zanamivir, the latter constitutes the alternative antiviral option against influenza A(H1N1)pdm09-H275Y variants. However, we and others have recently shown that the switch to zanamivir could induce zanamivir resistance mediated by substitutions at codon 119 (E119A/D/G) within the framework of the NA enzyme [11-14]. Noteworthy, A(H1N1)pdm09 variants with the double H275Y-E119D/G substitution are resistant to all available NAIs [11, 12].

Herein, we report a clinical case of multidrug resistance in an HSCT recipient patient who received oseltamivir and zanamivir therapies. By analysing the evolution of surface (HA and NA) genes from sequential respiratory samples, we confirmed the molecular pattern leading to the cross-resistance phenotype, as suggested by previous reports, and highlighted the potential compensatory role of HA mutations within the receptor binding site.

## 2. Materials and Methods

### 2.1. Molecular detection of influenza A(H1N1)pdm09 virus and antiviral treatment

A total of 11 nasopharyngeal swabs (NPS) were collected from an immunocompromised patient between 2018-01-27 and 2018-04-20 (Table 1). During that time, the patient received two courses of oseltamivir (from 2018-01-28 to 2018-02-14 and from 2018-02-27 to 2018-03-1) and one course of zanamivir (from 2018-03-01 to 2018-03-20). Plasma samples were also collected at some of these dates. RNA was extracted from clinical samples using the QIAsymphony SP automated instrument and served for a multiplex real-time PCR detection of respiratory pathogens using the 21 Fast Track Diagnostics system [15].

**Table 1.**
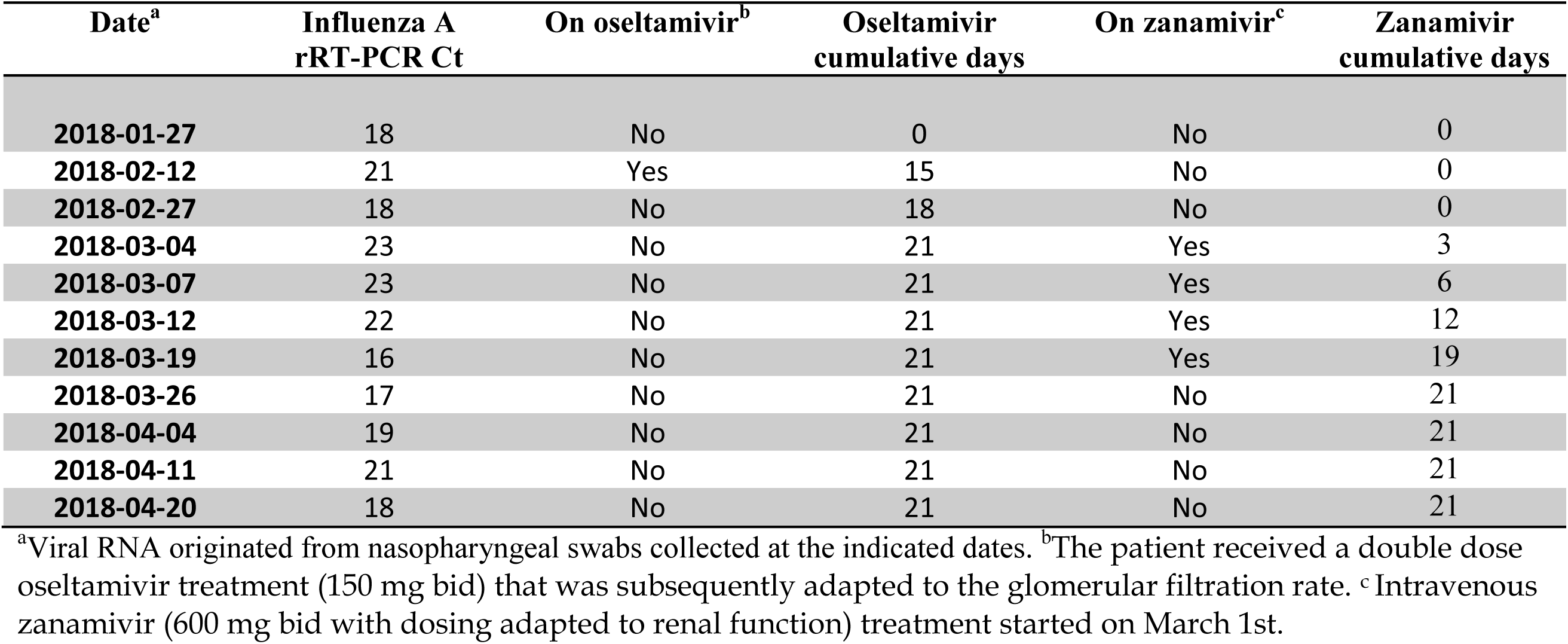
Molecular detection and NAI therapies during influenza A(H1N1)pdm09 infection in an immunocompromised patient.

### 2.2. Sequencing of viral surface genes

The HA and NA genes from nasopharyngeal (NP) swab samples of A(H1N1)pdm09 viruses were amplified by gene-specific RT-PCR tests and sequenced using the ABI 3730 DNA Analyzer (Applied Biosystems, Carlsbad, CA). The deduced amino acid sequences were compared after an alignment using the ClustalW program.

### 2.3. Viral quantification using droplet digital PCR (RT-ddPCR)

Viral RNA was extracted from thawed specimens using the QIAmp Viral RNA Mini Kit (QIAGEN, Mississauga, Ontario). The ddPCR workflow and data analyses were performed with the OneStep RT-ddPCR supermix according to the manufacturer’s instructions. The primers and probes targeting the H275 (WT) and Y275 (mut) variants of influenza A(H1N1)pdm09 virus were previously described [16]. The cycling and post-cycling steps were done as previously reported [17]. The cycled plate was then transferred and read in the FAM and HEX channels of the QX200 reader (Bio-Rad).

### 2.4. Generation of 7:1 reassortant influenza A(H1N1)pdm09 viruses

The pre-therapy and post-zanamivir HA genes from respiratory swabs were amplified by RT-PCR and cloned into the bidirectional pLLB-A plasmid as previously described [18]. Reverse genetics was used to generate 7:1 reassortant influenza viruses in the background of A/Québec/144147/09 (an A(H1N1)pdm09-like virus; GenBank accession numbers FN434457-FN434464) as previously reported [19].

### 2.5. Hemagglutination and elution assays

Hemagglutination assays were performed in U-bottom microtiter plates using 50 μL of serial 2-fold dilutions of 7:1 reassortant (generated as described above) and 50 μL of a 1% suspension of guinea pig or chicken red blood cells (RBCs) containing high proportions of α2,6-sialic acid (SA) receptors or α2,3-SA receptors, respectively. Plates were incubated at 4 °C for 1 h for determination of hemagglutination titers. For elution experiments, an equivalent of 16 HA units of each isolate were used for hemagglutination assays as stated above and the plates were transferred to 37 °C for recording elution times.

### 2.6. Patient description

A 72-year old man with acute myeloid leukemia underwent an allogenic HSCT on December 6, 2017. He was hospitalised on January 22, 2018, for suspected intestinal graft versus host disease (GvHD). Five days after admission, the patient presented influenza-like symptoms and a real-time RT-PCR test, performed on a nasopharyngeal swab (NPS), was positive for influenza A(H1N1)pdm09 virus. Double dose oseltamivir treatment [150 mg bid, subsequently adapted to the patient’s glomerular filtration rate (GFR)] was initiated on January 28. A chest computerized tomography (CT) scan performed the following day revealed the presence of bilateral ground glass infiltrates. Despite viral positivity on a NPS collected on February 12 (Table 1), oseltamivir was stopped on February 14, 2018, after 18 days of treatment, due to further decline in renal function. At the respiratory level, the patient’s clinical condition was stable at that time. However, oxygen saturation subsequently decreased again, and a chest CT scan performed on February 27 showed an increase of the pulmonary infiltrates. A NPS was collected and oseltamivir was re-introduced on February 27 (the dosage was adapted to a decreased GFR). In the absence of clinical improvement, intravenous (iv) zanamivir was started on March 1st, replacing oseltamivir. While subsequent influenza A RT-PCR cycle threshold (Ct) values initially suggested a decrease in viral load in NPS samples, the Ct values dropped again after March 12, despite continuous iv zanamivir (Table 1). However, the patient improved clinically, and a chest CT scan performed on March 20, showed a regression of the infiltrates, and zanamivir was therefore stopped. This exam also revealed an increase in bronchiectasis, indirectly supporting a fibrosing process, possibly due to pulmonary GvHD. On April 20, the patient experienced a novel episode of respiratory deterioration, and a chest CT scan performed on the same day showed a new increase of the ground glass infiltrates. Zanamivir was temporarily introduced again on April 20 but, due to a rapid clinical improvement upon broad spectrum antibiotic treatment initiation, antiviral treatment was interrupted after three days. Of note, all NPS samples collected from March 19 to April 20 were influenza A positive (Table 1). Thereafter, the patient became dyspneic again, and a chest CT scan performed on May 3rd displayed an increase in the pulmonary infiltrates. The patient’s condition deteriorated, particularly at the respiratory level, and he died on May 23rd. Autopsy revealed multiple pneumonia foci, of different ages, with active abscess forming pneumonia, organizing pneumonia foci, and constituted fibrosis.

## 3. Results

### 3.1. Molecular detection of influenza A(H1N1)pdm09 virus in clinical respiratory samples

The real time PCR for influenza A was positive for the 11 tested NPS samples with Ct values ranging between 16 and 23 (Table 1). Plasma samples collected on 2018-02-25 and 2018-02-26 were also positive (Ct of 37 and 38, respectively) whereas 4 other ones (collected on 2018-01-22, 2018-01-29, 2018-02-14 and 2018-03-05) were negative (not shown). Further sequencing experiments identified the influenza A viruses as A(H1N1)pdm09 variants and the sequences of NA and HA genes were determined.

### 3.2. Analysis of the NA genes

The focus was initially on the detection of H275Y substitution responsible for resistance to oseltamivir. In addition to Sanger sequencing that demonstrated the presence of mixed 275H/275Y chromatogram peaks, dd-RTPCR tests were performed for further quantification of 275Y and 275H viral populations. As summarized in Figure 1, the H275Y substitution could be detected as a major population (98.33% vs the WT) in the 2018-02-12 sample (collected after the first course of oseltamivir therapy). After a relative drop to 30.63% in the sample of 2018-02-27 (13 days after cessation of the first oseltamivir treatment), the variant persisted in the remaining post oseltamivir and post-zanamivir samples at relatively important rates (between 64.3% for sample 2018-03-19 to 99.64% for sample 2018-04-11). In addition to the H275Y substitution, the NA protein of samples collected during the first course of zanamivir therapy contained mixed residues at positions 287 (E/K), 94 (V/I), 95 (S/G) and 119 (E/G/D). Finally, the NA of the last three samples collected in the absence of therapy harbored a H275Y/E119G variant (Figure 1).

**Fig.1.**
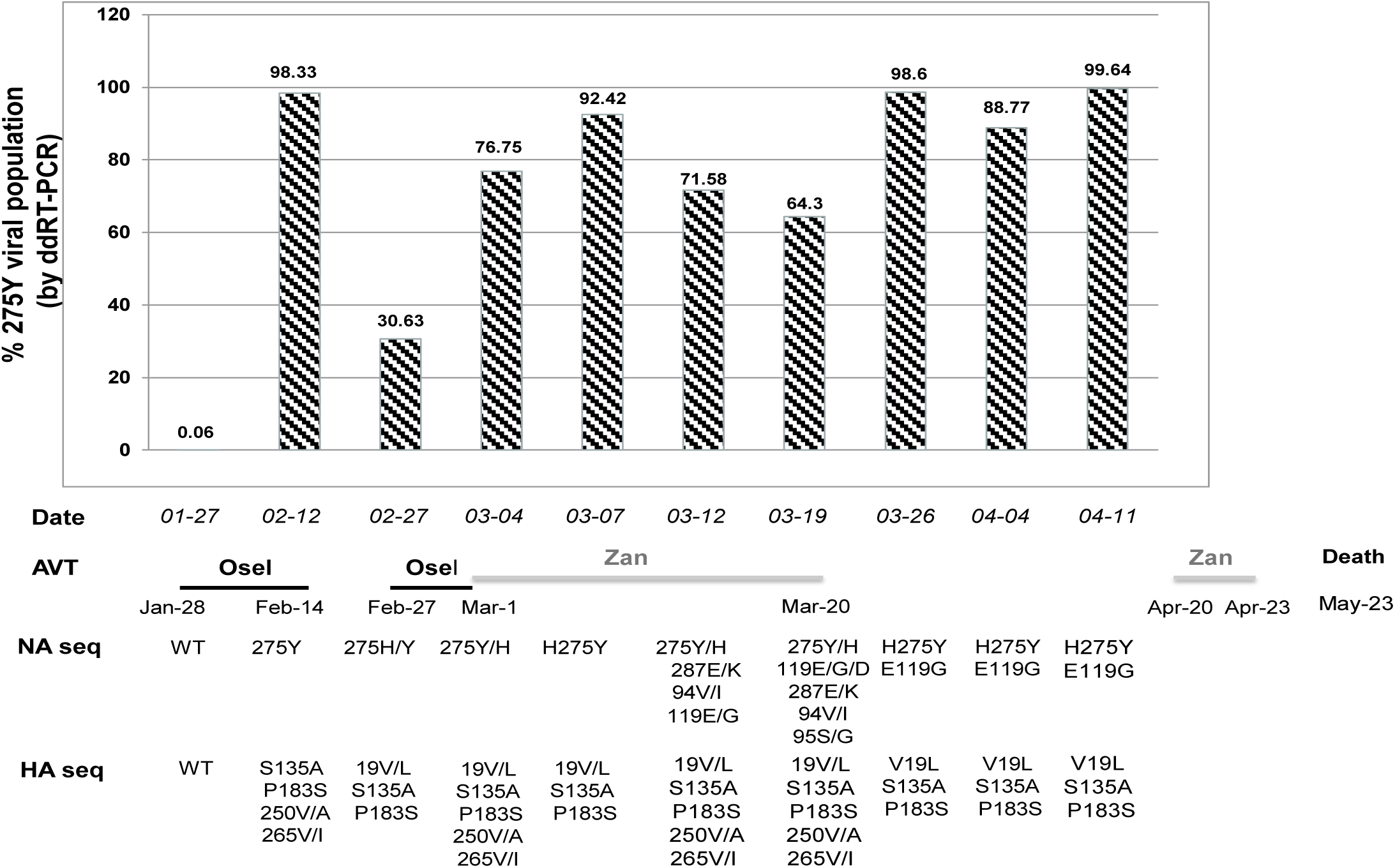
Molecular characterization of hemagglutinin (HA) and neuraminidase (NA) genes of influenza A(H1N1)pdm09 strains from an immunocompromised patient who received oseltamivir and zanamivir therapies. Characterization included ddRT-PCR quantification of NA 275H/Y populations (N1 numbering). The bars represent the % of 275Y population during oseltamivir (Osel) and zanamivir (Zan) therapies. Main NA and HA substitutions (H1 numbering, starting after the signal peptide) identified by Sanger sequencing are shown (bottom).

While the impact of the framework H275Y and E119G substitutions on the resistance phenotype to NAI is well documented [12], the effect of the remaining NA substitutions at residues 94, 95 and 287 has not been reported. We found no impact of these residues on NA expression and/or activity using recombinant proteins [20] (not shown).

### 3.3. Analysis of the HA genes

The HA gene of clinical samples contained three substitutions (V19L, S135A and P183S) that appeared in NPS collected after the first oseltamivir course and were subsequently maintained. Changes at residues 250 (V/A) and 265 (V/I) could also be transiently detected. The alignment of HA amino acid sequences (Figure 2) showed that the S135A and P183S substitutions were located near the receptor binding site of the HA molecule. Residue 135 is part of the 130-loop whereas residue 183 is near the 190-helix.

**Fig2.**
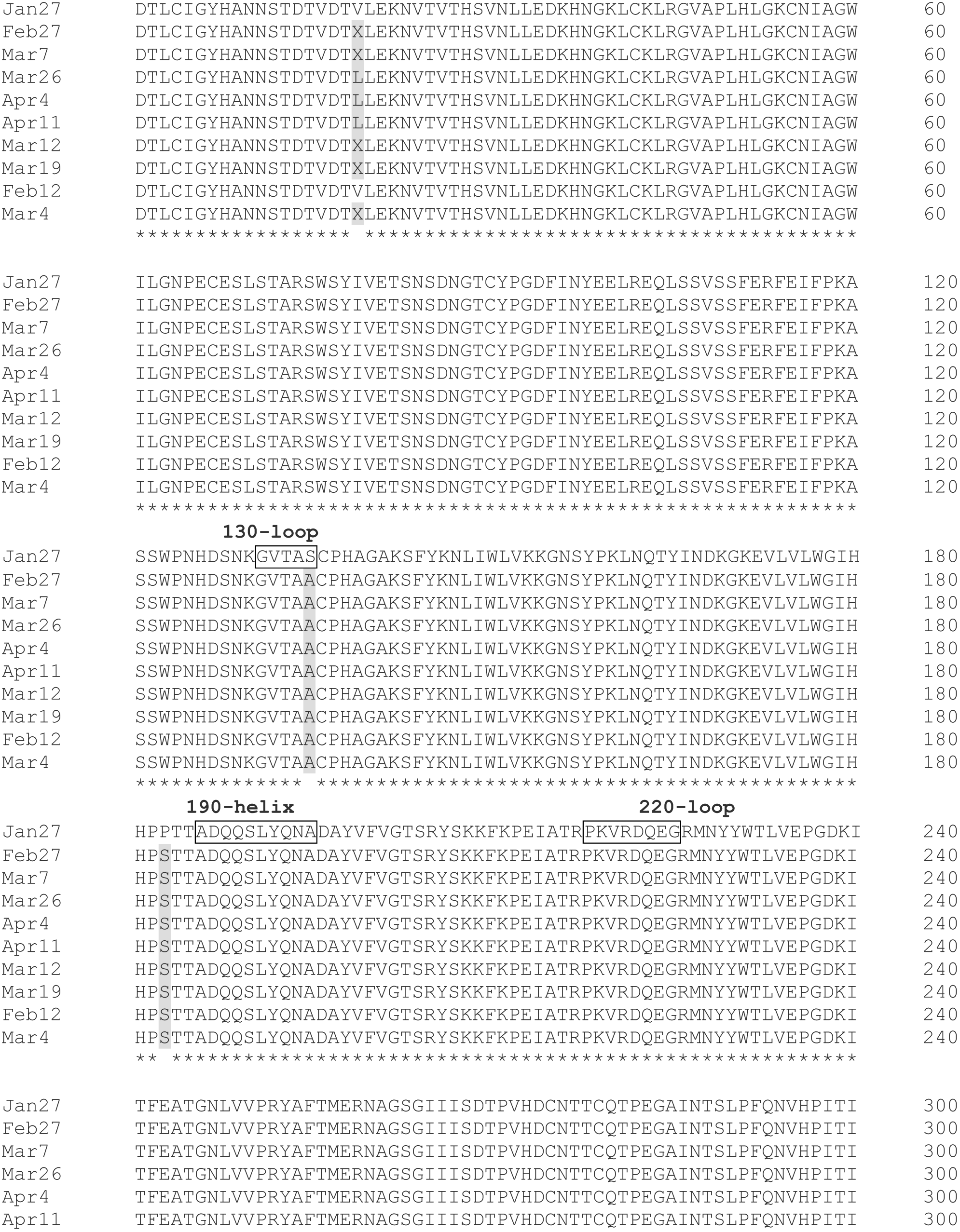
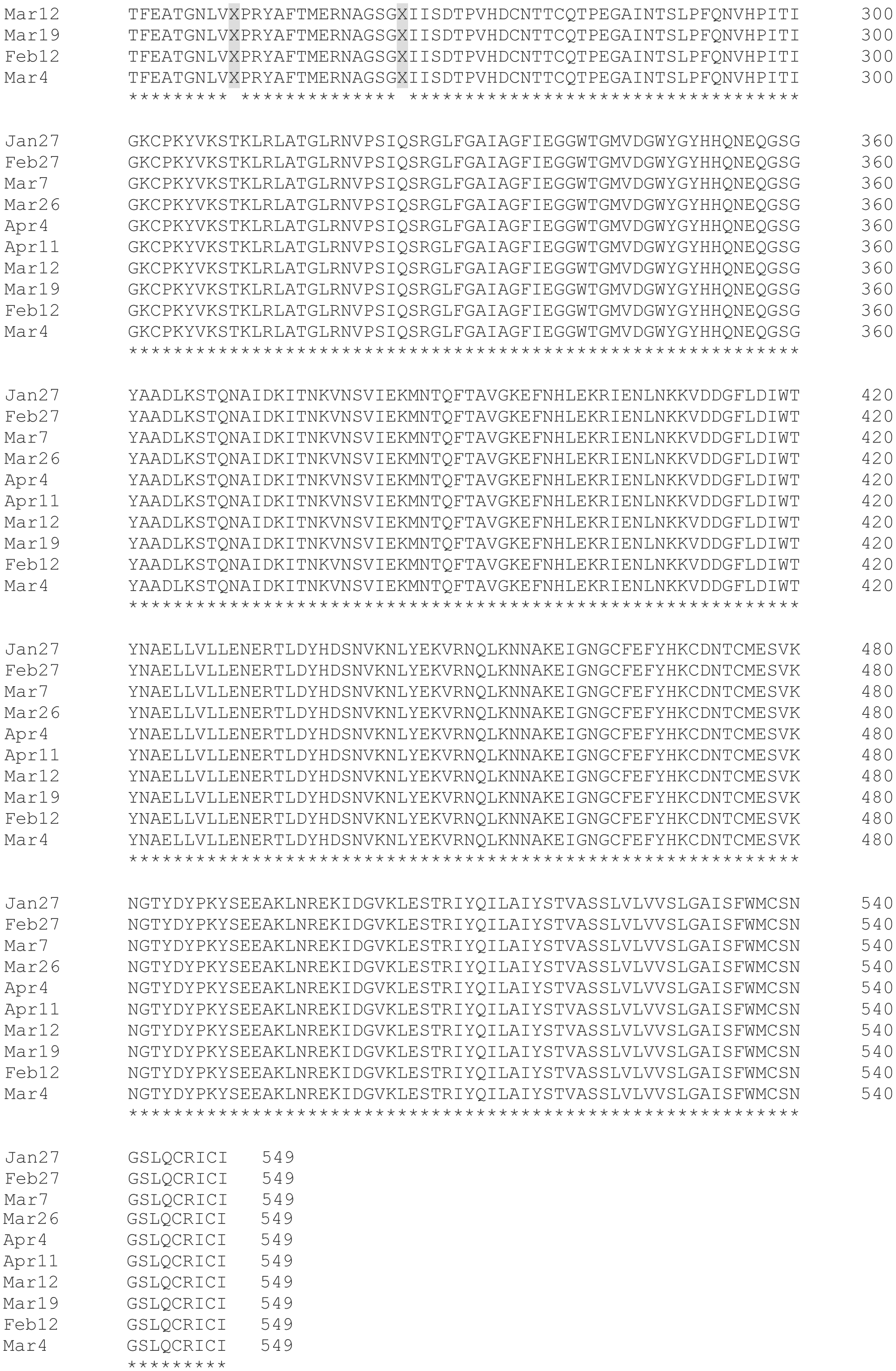
Evolution of the hemagglutinin (HA) of influenza A(H1N1)pdm09 strains from an immunocompromised patient who received oseltamivir and zanamivir therapies. Sequence analysis was performed using the CLUSTAL multiple alignment of the HA amino acid sequences. Sequences included that of the pre-therapy sample, collected on 2018-01-27 (Jan 27) and those from the post-NAI samples described in Table 1. Regions involved in the HA receptor binding site are indicated in boxes. Residues that differed versus the pre-therapy sequence are shaded. The X letter denotes mixed residues whereas asterisks denote identical residues.

In hemagglutination assays using A(H1N1)pdm09 7:1 reassortants, those containing the post-treatment HA (19L, 135A and 183S) eluted more rapidly guinea pigs red blood cells than the reassortants containing the pre-therapy HA (19V, 135S and 183P) (1 h vs > 4 h) (Table 2). The two reassortants had the same elution time (1 h) when using chicken red blood cells.

**Table 2.** Receptor binding specificities of A(H1N1) viruses using hemagglutination (HA) and HA elution assays.

| 7:1 reassortant<br>A(H1N1)pdm09<br>viruses | HA titer<sup>a</sup> |  | Elution time/gp-RB<sup>b</sup> | Elution time/cRBC<sup>b</sup> |
| --- | --- | --- | --- | --- |
| | gpRBCs<br>(α2,6 Gal) | cRBCs<br>(α2,3 Gal) | | |
| **HA: 19V, 135S, 183P**<br>**(Pre-therapy)** | 64 | 32 | > 4 h | 1 h |
| **HA: 19L, 135A, 183S**<br>**(Post-therapy)** | 32 | 32 | 1 h | 1 h |

<sup>a</sup>Hemagglutination (HA) assays were performed using guinea pig or chicken red blood cells (gpRBCs and cRBCs, respectively). Binding was performed at 4 °C for 1 h. <sup>b</sup>For elution assays, U-bottom microwell plates containing 16 HA units of each virus were incubated at 37 °C for determining the time (in hours) required for completed viral elution.

## 4. Discussion

Reduced inhibition to NAIs in influenza viruses is mainly mediated by changes (substitution/deletions) in or near the active site of the NA enzyme [21]. Although NAIs are analogues of the sialic acid (DANA: 2,3-dehydro-2-deoxy-N-acetylneuraminic acid), zanamivir and oseltamivir can be distinguished by important structural differences that may influence the nature and rate of mutations of resistance [22]. Zanamivir, which is structurally closer to DANA compared to oseltamivir [23], has been infrequently associated with the emergence of viral resistance [6]. Accordingly, *in vitro* passaging of influenza A(H1N1)pdm09 virus under zanamivir pressure failed to induce drug resistance [24]. On the other hand, in A(H1N1)pdm09 infections, prophylactic or therapeutic exposures to oseltamivir readily selected for NAI-resistant H275Y variants in immunocompromised patients [7, 11, 12]. Besides NA changes, reduced inhibition to NAIs could also involve changes in or near the receptor binding site (RBS) of the HA protein due to the important functional balance between HA and NA surface proteins [25]. The RBS of influenza viruses consists of three structural elements including two loops (130-loop and 220-loop) and one helix (190-helix) [26].

In this, report, we took advantage of having a series of respiratory samples from an HSCT patient including pre-therapy NPS and samples collected after oseltamivir (used as a first-line antiviral agent) and zanamivir (administered after detection and persistence of the H275Y substitution) for analyzing the evolution of influenza A(H1N1)pdm09 NA and HA genes. Similarly to previous reports, we found that oseltamivir rapidly selected for the H275Y NA mutation (Figure 1). In fact, a very low proportion of that variant (0.06%) could be detected by ddRT-PCR in the first NPS collected before oseltamivir therapy. Oseltamivir pressure enhanced the H275Y level to 98.3 % in the subsequent NPS. As previously described [11-13], the switch to zanamivir also induced the occurrence of a second NA substitution at the framework E119 residue (E119G). This substitution was maintained during one month in the double H275Y-E119G variant (Figure 1). Tamura and colleagues reported that the dual H275Y-E119G mutation recovered from an immunocompromised child resulted in ≥100-fold reduced inhibition against zanamivir, oseltamivir, peramivir and laninamivir [12]. This was also the case for another H275Y-E119D A(H1N1)pdm09 variant described by us [11]. Of interest, viral cultures could not be obtained for the two double mutants suggesting attenuating effects [11, 12]. The role of the H275Y-E119D change in altering viral fitness was further confirmed both *in vitro* and in animal models [20].

As shown in Figure 1, besides NA changes, the clinical A(H1N1)pdm09 variants contained HA substitutions under NAI pressure. As S135A and P183S changes are located in or near the HA RBS, there was a possibility for altered HA binding property. For instance, a recent report showed that *in vitro* passaging of A/California/04/2009 (H1N1) virus under lectins pressure induced the S183P HA substitution, which significantly increased binding to α2,6 Sa-linked glycans [27]. In a recent clinical report, NAI therapy against A(H1N1)pdm09 infection in a child with severe combined immunodeficiency disease (SCID) induced the P183S HA change [14]. In our HA elution experiments, the reassortant containing the pre-therapy HA (135S, 183P) had a higher affinity to the guinea pigs RBC (rich in α2,6 Sa-linkage) from which it could not elute after 4 h of incubation at 37 °C contrasting with the complete elution within 1 h for the reassortant containing the post-therapy (183S) HA (Table 2). Of note, while most influenza H1N1 strains of 2018 harbored HA with 135S and 183P residues, the HA of previous 2009 and 2015 vaccine strains (A/California/07/2009 and A/Michigan/45/2015, respectively) had A and S at these positions, respectively (Table S). Therefore, the NAI pressure seemed to cause a reversion to the original A(H1N1)pdm09 genotype. Thus, due to the reduced NA activity caused by the H275Y substitution, the isolate had to select for HA substitutions with the potential to reduce affinity to α2,6 receptors for restoring the HA/NA balance, as demonstrated in our HA elution experiments. Unfortunately, the fact that we do not possess the clinical isolates with the different HA/NA genotypes to compare their fitness constitutes a limitation of this study. Moreover, we could not rescue recombinant viruses harboring the post-therapy NA protein (H275Y/E119G) despite many attempts. Finally, the potential compensatory role of changes in other viral segments is also to be considered.

In conclusion, we provide herein an additional clinical report on the emergence of multidrug-resistant A(H1N1)pdm09 viruses in immunocompromised patients receiving sequential oseltamivir and zanamivir therapies. In conjunction with previous reports, there seems to be a consistent molecular pathway leading to the multi-drug resistance phenotype (Figure S). It consists on the rapid selection of the H275Y NA substitution, following oseltamivir therapy, followed by E119G/A/D changes while on zanamivir therapy. The resulting double NA mutant displays highly reduced susceptibility to all NAIs. Contrasting to previous reports [11-13], we provide here insights for possible involvement of HA changes in or near the RBS on the drug resistance phenotype. Due to limited antiviral options for the control of influenza infections in immunocompromised individuals, the occurrence of such molecular events may have major clinical implications. This report highlights the need for the development of novel anti-influenza compounds

## 5. Acknowledgements

This work was supported by a Canadian Institutes of Health Research (CIHR) foundation grant to GB (grant No. 229733) for a research program on the pathogenesis, treatment and prevention of respiratory and herpes viruses.

## Funding

No private funding

## Competing interests

None declared

## Ethical Approval

Not required

## Legends

**Figure S.**

**Schematic representation of the molecular pathway leading to the emergence of multi-drug resistant A(H1N1)pdm09 variants in immunocompromised patients.** The pathway is based on this study and recent clinical reports by L’huillier et al. [11], Tamura et al. [12], Trebbien et al. [13] and Pichon et al. [14]. The pathway suggests that the use of oseltamivir as first-line antiviral agent induces the H275Y NA substitution, responsible for resistance to oseltamivir and peramivir. The subsequent switch to zanimivir induces changes at residue E119 conferring resistance to all NAIs. HA substitutions within the receptor binding site could be also selected for potentially restoring a functional HA/NA balance. Note : Osel, oseltamivir ; Zan, zanamivir ; Per, peramivir ; Lan, laninamivir ; RBS, receptor binding site, NA neuraminidase.

